# Enhanced keratin extraction from wool waste using a deep eutectic solvent

**DOI:** 10.1101/2021.09.29.462276

**Authors:** Oseweuba Valentine Okoro, Hafez Jafari, Parinaz Hobbi, Lei Nie, Houman Alimoradi, Amin Shavandi

## Abstract

In this study, the solubilisation of waste coarse wool as a precursory step for the large scale valorisation of keratin was investigated using a green deep eutectic solvent (DES) based on L-cysteine and lactic acid. The investigation was undertaken via the response surface methodology and based on the Box-Behnken design for four process variables of temperature (70-110 °C), dissolution time (2-10 h), the mass of L-cysteine (0.5-2.5 g) in 20 mL of lactic acid, and wool load in the DES (0.2-0.6 g). The effect of variations in temperature was established to be the most significant process variable influencing keratin yield from waste coarse wool in the current work. An optimum keratin yield (93.77 wt.%) was obtained at the temperature of 105 °C, 8 h dissolution time, with 1.6 g L-cysteine in 20 mL of lactic acid using 0.5 g of wool. This study suggests L-cysteine and lactic acid as a green solvent with the potential to scale up keratin recovery from waste wool without significant destruction in the structure of the recovered keratin.

**Highlights:**

- Keratin recovery from wool using deep eutectic solvent was assessed
- The basis for the use of the new deep eutectic solvent was discussed
- The effects of the process variables on keratin yield were explored
- Keratin recovered was optimised and characterised.

## 1 Introduction

Keratin, the third most abundant polymer in nature after chitin and cellulose, is a renewable, biocompatible, and biodegradable fibrous protein present in hair, coarse wool, feathers, nails, and horns [1–3]. Keratin can exist structurally as α-helix structures (α-helix keratins) and ß-sheet structures (β-keratins) [4, 5]. Alpha (α)-helix keratins are present in vertebrates and specifically in hooves and coarse wool, while β-keratins are present in the epidermal appendages of reptiles and birds [4, 5]. Keratin has a molecular weight ranging from ~40 kDa to ~68 kDa [6] and contains cysteine (~7 wt.% to 20 wt.%) [1]. The presence of cysteine facilitates the formation of strong inter-and intra-molecular disulfide bonds [1]. Keratin is therefore insoluble in most solvents which makes its dissolution and regeneration challenging [7]. Notably, keratin has a complex hierarchical-like filament-matrix structure containing carboxylic functional groups and polypeptide chains that make it chemically reactive when the functional groups are ‘exposed’ via reduction or protonation of disulfide cross-links [8, 9]. Due to the high chemical activity and biocompatibility of keratin, it can be employed in the manufacture of several value-added products such as bioadhesives and fibrous composite materials [9, 10]. Additionally, keratin can be used in the development of biomedical gels, films, nano/micro-particles, and beads. [9] Modified keratin can also serve as a bio-sorbent for removing toxic metal ions from water resources on account of the exceptionally important role of functional groups for keratin-metal binding [9]. Given the economic and environmental benefits of value extraction from waste wool, keratin recovery has been widely studied in the literature [11–15]. To facilitate keratin extraction from waste coarse wool, it is necessary to split the intra- and intermolecular disulfide and hydrogen bonds [16]. Several methods have therefore been explored for keratin extraction from waste coarse wool via protein denaturation using reducing agents and hydrogen bonding disrupters such mercaptoethanol and urea/ thiourea, and urea/metabisulfite respectively [17, 18]. The use of these chemicals however present associated issues of toxicity, high cost and may lead to the production of a keratin product that is characterized by low molecular weight fragments [17–19]. Another approach for the dissolution of waste wool is the utilization of deep eutectic solvents (DESs) which are considered green, simple, and effective extraction solvents. DESs consist of a mixture of compounds acting as hydrogen bond acceptor and hydrogen bond donors, thus are capable of forming intermolecular hydrogen bonds and Van der Waals interactions with the wool structure [20]. For instance, in a previous study the utilization of a deep eutectic solvent mixture of choline chloride/oxalic acid was employed for keratin extraction from waste wool [21]. In that study, 5 wt. % of the wool was shown to be completely dissolved in the DES at high temperatures of up to 125 °C [21]. We recently developed a new, efficient and biocompatible DES composed of lactic acid and L-cysteine for wool dissolution [22]. This study [22] established the functionality of lactic acid and L-cysteine utilization in the production of a nontoxic and biodegradable DES that can facilitate significant keratin recovery from wool without compromising the keratin structure [22]. Indeed a previous study has shown that the use of this novel DES has the potential of producing keratin composed of proteins with high molecular masses [22]. In the current study, we are therefore seeking to optimize the recovery of keratin from waste coarse wool using this new DES composed of lactic acid and L-cysteine. We are also seeking to investigate the physicochemical properties of the keratin obtained at different processing conditions.

## 2 Material and methods

### 2.1 Materials and chemical reagents

Raw waste coarse wool, free from coarse fertilizer and litter, was locally sourced. The coarse wool was defatted via Soxhlet extraction using the solvent mixture of 1:1 v/v hexane and dichloromethane. The defatted coarse wool was then initially rinsed with distilled water and subsequently dried in an oven at 60 °C to constant mass. The wool was then stored in a clean polythene bag at room temperature before undertaking the experimental runs. Dialysis membrane with 500 Da molecular weight cut-off and all analytical grade solvents and reagents used in this study (unless otherwise stated) were obtained from Fischer scientific (Merelbeke, Belgium). The chemical reagents of L-cysteine (analytical reagent) and lactic acid (analytical reagent) were purchased from Merck (Darmstadt, Germany) and Fischer scientific (Merelbeke, Belgium), respectively. Hexane, dichloromethane and the dialysis membrane (with 500 Da molecular weight cut-off) were purchased from Fischer scientific (Merelbeke, Belgium).

### 2.2 Keratin extraction from wool

According to the experimental design, in each experimental run, a mass of the dried-defatted coarse wool was treated with the DES, containing a known mass of L-cysteine in 20 mL of concentrated lactic acid, at a specific temperature and dissolution time. The ranges of the temperature, dissolution time, mass of L-cysteine and wool load investigated were specified as 70-110 °C, 2-10 h, 0.5-2.5 g and 0.2-0.6 g, respectively. The wool solubilisation was undertaken in a reactor equipped with a magnetic stirrer for uniform mixing of the reaction components with heating achieved using a glycerol bath. The reactor temperature was maintained using a proportionalintegral-derivative (PID) temperature controller. The upper and lower limits of the processing parameters were selected in accordance with our preliminary results and the reported values in the literature. [11, 23, 24] In particular, due to concerns regarding keratin degradation at high temperatures (i.e. > 130 °C [11]), the current study specified the upper limit of the temperature as 110 °C. At the end of each experiment, the resulting mixture was dialyzed using a 500 Da membrane against distilled water for 3 days at room temperature, and the water was replaced every 8 h. The recovery of the keratin product was achieved via freeze-drying of the mixture, after which the mass of the resulting keratin powder was measured, and the keratin yield in percentage (*K_y_*) calculated as follows;

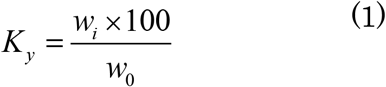

where, *w_0_* and *w_i_* denote the weight of initial coarse wool fiber and the weight of recovered keratin powder in g, respectively.

### 2.3 Experimental design and statistical analysis

The Box-Behnken experimental design (BBD) was used in the present study since it is recognized as highly proficient and low cost to execute. [25, 26] The present study seeks to investigate the influence of selected process variables of mass of L-cysteine, (*mL-cys*), reaction temperature in °C (*T*), coarse wool dissolution time in *h* (*t*) and coarse wool load in g (*W*) on the keratin yield. Having specified the ranges of the process variables, these values were subsequently coded to aid statistical calculations as follows [27];

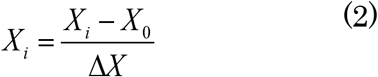

where X_i_ denotes the coded value of the process variable; X_i_ is the process variable’s actual value; X_0_ denotes the actual value of X_i_ at the center point with the step change value denoted as ΔX.

The statistical analysis of the experimental results was performed using Minitab^®^ 17.1.0 (Minitab, Inc. USA). The study was achieved by undertaking 27 experimental runs. Based on the data from the Box-Behnken experimental design, an empirical relation between keratin yield and the process variables was developed. The generated function was of the second-order polynomial form as follows;

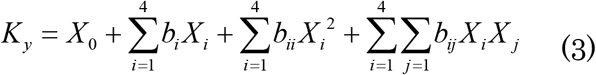

where *K_y_* denotes the keratin yield (wt. %), *X_o_* represents the model intercept, *X_i_* (*X_j_*) represents the *i*th (*j*th) system variable (temperature, dissolution time, mass of L-cysteine in 20 mL of lactic acid, or coarse wool load), *b_i_, b_ii_* and *b_ij_* represent the model regression coefficients.

Table 1 summarizes the values of the process variables investigated and their associated coded levels employed in the Box–Behnken design method.

**Table 1.** Coded and actual levels utilized in the Box-Behnken experimental method.

| Parameters | Coded and actual values for the levels in the experimental design |  |  |
| --- | --- | --- | --- |
|  | low | centre | high |
| Levels | -1 | 0 | +1 |
| Mass of L-cysteine, $m_{L-cys}$ (g) | 0.5 | 1.5 | 2.5 |
| Dissolution time, $t$ (h) | 2 | 6 | 10 |
| Temperature, $T$, (°C) | 70 | 90 | 110 |
| Coarse wool load, $W$ (g) | 0.2 | 0.4 | 0.6 |

Based on the values of the levels specified in Table 1, different combinations of the process variables were subsequently assessed according to the experimental design and are presented in the results and discussion section.

Furthermore, the significance of each process variable on the keratin yield was assessed via the consideration of the student *F*-value of each variable relative to the critical *F*-value of the experimental data and design. This approach is well-known and has been utilized in the literature[28–30]. Based on this approach, the more the student *F*-value of the process parameter exceeds the critical *F*-value, the more significant is the effect of variations in the process variable on the keratin yield. The critical *F*-value of the present work was based on the experimental design and was determined to be 3.13. For completeness, the conditions of the process variables that facilitate an optimal keratin yield were determined using the desirability function algorithm in Minitab. Discussions related to describing the theoretical basis of the desirability function are beyond the scope of the present work and are presented elsewhere [31]. After wool solubilisation was achieved, the filtrate containing the dissolved wool was dried to constant mass for keratin recovery. The dried keratin was then collected, sealed in an airtight bag and stored in a fridge at the temperature of 4 °C before undertaking characterization experiments.

### 2.4 Characterization experiments

Dried keratin samples were selected and their physiochemical properties, subsequently investigated. Investigations undertaken included the Fourier-transform infrared spectroscopy (FTIR), differential scanning calorimetry (DSC), thermogravimetric analysis (TGA) and X-ray diffraction (XRD) analyses. The FTIR investigation was undertaken using the Infrared Spectrometer (Jasco brand model FT/IR-6600, equipped with Spectra Manager™ II Software) in a wavelength range of 4000 to 400 cm^-1^, with a 4 cm^-1^ spectral resolution at room temperature. The DSC assessment of the keratin sample was performed using a Mettler Toledo DSC 821. The analysis was performed under a dynamic nitrogen atmosphere (20 mL/min) using a sample mass of 10 ± 0.1 mg and a heating rate of 10°C/min. The TGA of samples was undertaken using a TA instrument (Netzsch STA 409 PC coupled with Netzsch QMS 403C). The dried keratin sample (50 mg) was heated from the temperature of 30 °C to 600 °C at 10 °C/min heating rate under nitrogen flow (60 mL/min). The XRD experiment employed a diffractometer (XRD; Bruker ecoD8 advance), operating in the range of 5°<2θ<60° with Cu Kα irradiation (k=0.154060 nm) at 40 kV and 25 mhA to generate XRD patterns. The diffractometer was operated at a scanning rate of 1.2°/min with a step size of 0.019 and scan step time of 96.00s [32].

## 3 Results and discussions

### 3.1 Keratin extraction

#### 3.1.1 Model fitting

The experimental results highlighting the keratin yields for different experimental conditions are presented in Table 2. The yield of keratin was found to range from 5.95 wt.% to 90.04 wt.%, indicating the impact of the processing conditions selected in the study. Based on the specified model (in Equation 3) and the resulting experimental data, the relation between keratin extraction yield and the mass of L-cysteine in 20 mL of lactic acid, *m_L-cys_*, reaction temperature *T*, coarse wool dissolution time *t* and coarse wool load, *W* was determined as follows;

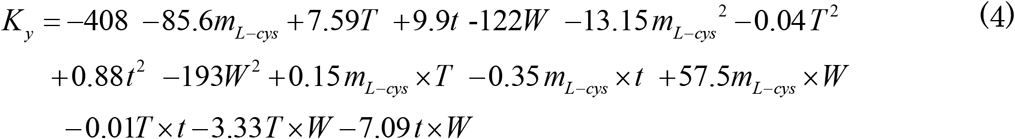

**Table 2.** Keratin yield at different process variable conditions.

| N | Coded values of the parameters |  |  |  | Actual values of the parameters |  |  |  | Response |
| --- | --- | --- | --- | --- | --- | --- | --- | --- | --- |
| | $m_{L-cys}$(g) | $T$(°C) | $t$(h) | $W$(g) | $m_{L-cys}$(g) | $T$(°C) | $t$(h) | $W$(g) | |
| 1 | 0 | -1 | 0 | 1 | 1.5 | 70 | 6 | 0.6 | 5.95 |
| 2 | 0 | -1 | -1 | 0 | 1.5 | 70 | 2 | 0.4 | 22.15 |
| 3 | -1 | -1 | 0 | 0 | 0.5 | 70 | 6 | 0.4 | 11.45 |
| 4 | -1 | 1 | 0 | 0 | 0.5 | 110 | 6 | 0.4 | 72.30 |
| 5 | 1 | 0 | 1 | 0 | 2.5 | 90 | 10 | 0.4 | 73.55 |
| 6 | -1 | 0 | 1 | 0 | 0.5 | 90 | 10 | 0.4 | 31.96 |
| 7 | 0 | 0 | 1 | -1 | 1.5 | 90 | 10 | 0.2 | 85.72 |
| 8 | 0 | 0 | 0 | 0 | 1.5 | 90 | 6 | 0.4 | 77.77 |
| 9 | 0 | -1 | 1 | 0 | 1.5 | 70 | 10 | 0.4 | 31.67 |
| 10 | 0 | -1 | 0 | -1 | 1.5 | 70 | 6 | 0.2 | 32.46 |
| 11 | 0 | 0 | 0 | 0 | 1.5 | 90 | 6 | 0.4 | 79.93 |
| 12 | 0 | 1 | 0 | 1 | 1.5 | 110 | 6 | 0.6 | 81.54 |
| 13 | 1 | 0 | -1 | 0 | 2.5 | 90 | 2 | 0.4 | 46.51 |
| 14 | 0 | 1 | 1 | 0 | 1.5 | 110 | 10 | 0.4 | 90.04 |
| 15 | -1 | 0 | 0 | -1 | 0.5 | 90 | 6 | 0.2 | 51.62 |
| 16 | 0 | 1 | 0 | 0 | 1.5 | 110 | 2 | 0.4 | 77.95 |
| 17 | 1 | -1 | 0 | 0 | 2.5 | 70 | 6 | 0.4 | 36.12 |
| 18 | 0 | 0 | 0 | 0 | 1.5 | 90 | 6 | 0.4 | 76.73 |
| 19 | 0 | 0 | -1 | 1 | 1.5 | 90 | 2 | 0.6 | 21.00 |
| 20 | 0 | 0 | 1 | 1 | 1.5 | 90 | 10 | 0.6 | 80.62 |
| 21 | 0 | 0 | -1 | -1 | 1.5 | 90 | 2 | 0.2 | 48.71 |
| 22 | -1 | 0 | -1 | 0 | 0.5 | 90 | 2 | 0.4 | 10.59 |
| 23 | -1 | 0 | 0 | 1 | 0.5 | 90 | 6 | 0.6 | 67.35 |
| 24 | 1 | 1 | 0 | 0 | 2.5 | 110 | 6 | 0.4 | 84.68 |
| 25 | 1 | 0 | 0 | -1 | 2.5 | 90 | 6 | 0.2 | 85.78 |
| 26 | 1 | 0 | 0 | 1 | 2.5 | 90 | 6 | 0.6 | 55.47 |
| 27 | 0 | 1 | 0 | -1 | 1.5 | 110 | 6 | 0.2 | 54.72 |

The resulting R^2^ value for the developed model was determined to be 0.8569, suggesting that the regression model was sufficient to describe the experimental results more so given that the R^2^ value was greater than the minimum acceptable R^2^ value of 0.7 in scientific works [33, 34].

#### 3.1.2 Assessment of model statistics

The sufficiency of the empirical model was analyzed with analysis of variance (ANOVA) and shown in Table 3. The *F*-value of the model is determined to be 5.13, implying that the model is statistically significant (Table 3). Variations in the temperature (*F*-value of 38.28), dissolution time (*F*-value of 10.29) and mass of L-cysteine (*F*-value of 6.94) present the most significant independent impacts on the keratin yield since in these cases, the *F*-values exceed the critical *F*-value of 3.13. The effect of variations in the coarse wool load on the keratin yield is shown to be statistically insignificant since its *F*-value of 0.82 is less than the critical *F*-value of 3.13. Based on the magnitude of the *F*-value for the temperature parameter of 38.28, changes in temperature are therefore established to constitute the most impactful process variable in the system. Furthermore, except for the effect of the interaction of temperature and coarse wool load (*F*-value > 3.13), the effects of the interactions of other process parameters on keratin yield are shown to be insignificant.

**Table 3.** Analysis of variance (ANOVA) for the empirical model of the keratin yield.

| Source | DF | Sum of squares | Mean square | $F$-Value | $p$-Value | Remarks |
| --- | --- | --- | --- | --- | --- | --- |
| Model | 14 | 16166.1 | 1154.72 | 5.13 | 0.004 | ** |
| $m_{L-cys}$ | 1 | 1560.4 | 1560.43 | 6.94 | 0.022 | ** |
| $T$ | 1 | 8609.6 | 8609.61 | 38.28 | 0.000 | ** |
| $t$ | 1 | 2314.4 | 2314.35 | 10.29 | 0.008 | ** |
| $W$ | 1 | 184.7 | 184.69 | 0.82 | 0.383 | * |
| $m_{L-cys}^2$ | 1 | 921.6 | 921.60 | 4.10 | 0.066 | ** |
| $T^2$ | 1 | 1441.4 | 1441.37 | 6.41 | 0.026 | ** |
| $t^2$ | 1 | 1046.9 | 1046.88 | 4.65 | 0.052 | ** |
| $W^2$ | 1 | 316.4 | 316.38 | 1.41 | 0.259 | * |
| $m_{L-cys} \times T$ | 1 | 37.8 | 37.76 | 0.17 | 0.689 | * |
| $m_{L-cys} \times t$ | 1 | 8.0 | 8.04 | 0.04 | 0.853 | * |
| $m_{L-cys} \times W$ | 1 | 529.9 | 529.92 | 2.36 | 0.151 | * |
| $T \times t$ | 1 | 1.7 | 1.65 | 0.01 | 0.933 | * |
| $T \times W$ | 1 | 711.1 | 711.10 | 3.16 | 0.101 | ** |
| $t \times W$ | 1 | 127.7 | 127.69 | 0.57 | 0.466 | * |

\*low significance, i.e. $F$ -value < 3.13, \*\*high significance, i.e. $F$ -value > 3.13, DF denotes degrees of freedom

### 3.2 Independent effects of variations of the process variables on keratin yield

Further assessments of the independent effects of the process variables were explored using 2D plots that were based on Equation 4 and are highlighted in Figure 1. Figure 1 highlights the model-derived mean independent effects of the process variables, in the absence of process variable interaction effects and is discussed below.

**Figure 1.**
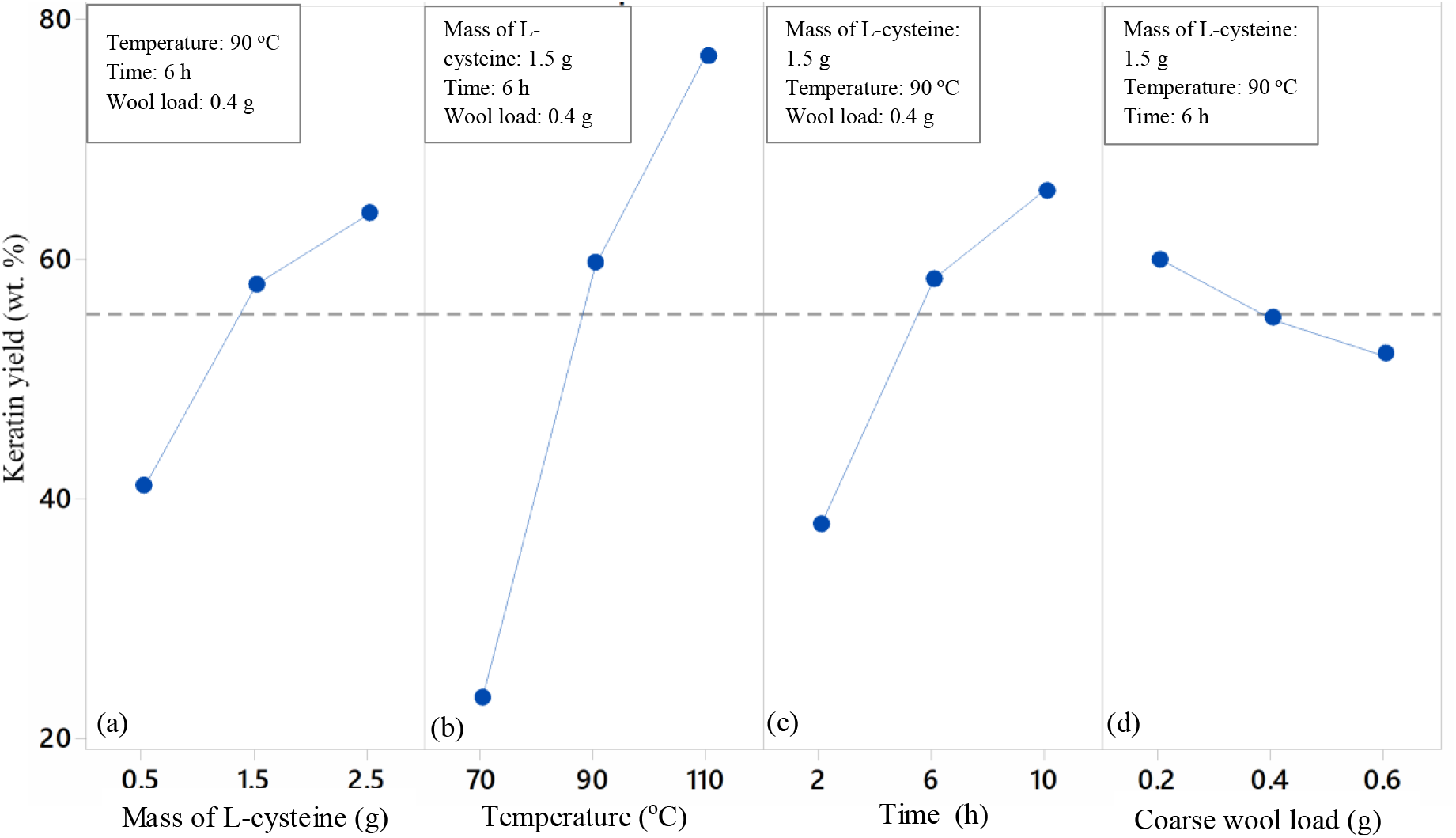
The independent effects of the process variables on keratin yield. (a) Denotes effect of the variation of mass of L-cysteine on keratin yield, (b) denotes the effect of the variation of temperature on keratin yield, (c) denotes the effect of the variation of time on keratin yield, (d) denotes the effect of the variation of coarse wool load on keratin yield. The legend specifies the conditions of the process parameters held constant.

#### 3.2.1 Effects of variations in the mass of L-cysteine

Figure 1(a) shows that as the mass of L-cysteine increases from 0.5 g to 2.5 g in 20 mL of lactic acid, the keratin yield increases from ~40 wt. % to ~65wt. %. This observation is expected when the role of L-cysteine in wool dissolution is understood. Initially, the lactic acid component of the DES compromises the bonds in the wool since lactic acid readily forms hydrogen bonds with peptides and other functional groups in wool.[20] The compromise of wool hydrogen bonds allows the cysteine to penetrate and cleave the disulphide bonds such that the increased the availability of L-cysteine facilitates the production of thiolate anions which are then available for the cleavage of disulphide bonds in keratin, leading to a higher wool solubilisation for keratin recovery

#### 3.2.2 Effects of variations in the temperature

As can be seen in Figure 1(b), the Keratin extraction yield was increased by heating the mixture (from 70 °C to 110 °C). The yield at 110 °C was over three times more than the yield at 70 °C (from 23 wt.% to 77 wt.%). This observation is due to the enhanced rate of the redox reactions involving L-cysteine and the disulphide bonds in keratin in the wool, as the temperature increases. The redox reactions involving L-cysteine facilitates the irreversible disulphide cleavage of keratin (R-S-S-R), leading to the formation of a disulfide bridge (R–S–S–SCH2CH(NH2)COO^-^). [35, 36] The rate of the redox reaction will increase as the temperature increases since higher temperatures lead to reduced viscosities for enhanced mobility and interaction of the thiolate anions from the cysteine and the disulphide bonds in keratin.

#### 3.2.3 Effect of variations in dissolution time

The effect of variations in the dissolution time, *t* in h on keratin yield is observed in Figure 1(c). Figure 1(c) shows that a longer dissolution time translates to a higher keratin yield. In Figure 1(c) increasing the dissolution time from 2 h to 10 h resulted in an average increase of keratin yield from 38 wt. % to 66 wt.%. This observation is consistent with our understanding of reaction kinetics, since longer dissolution times enhance opportunities for disulphide cleavages which are necessary for wool keratin dissolution.

#### 3.2.4 Effects of changes in coarse wool load

The effect of changes in coarse wool load in the DES in g on keratin yield is aptly highlighted in Figures 1(d). Figure 1(d) shows that the increase in the coarse wool load from 0.2 g to 0.6 g leads to a decrease in the keratin yield from 60 wt. % to 52 wt.%. This observation may be due to higher wool loads inhibiting the capacity of the lactic acid component of the DES to facilitate wool “swelling” via the compromise of wool bonds for L-cysteine enabled irreversible disulphide cleavage of keratin, for enhanced yields. This observation reinforces the need to investigate the preferred condition of the process variable of the wool load that will facilitate an optimum keratin recovery. Having explored the independent effects of the process variables, the combined effects, arising from the interaction of the process parameters was assessed using Equation 4. The combined effects were assessed using three dimensional surface plots in Figure 2.

**Figure 2.**
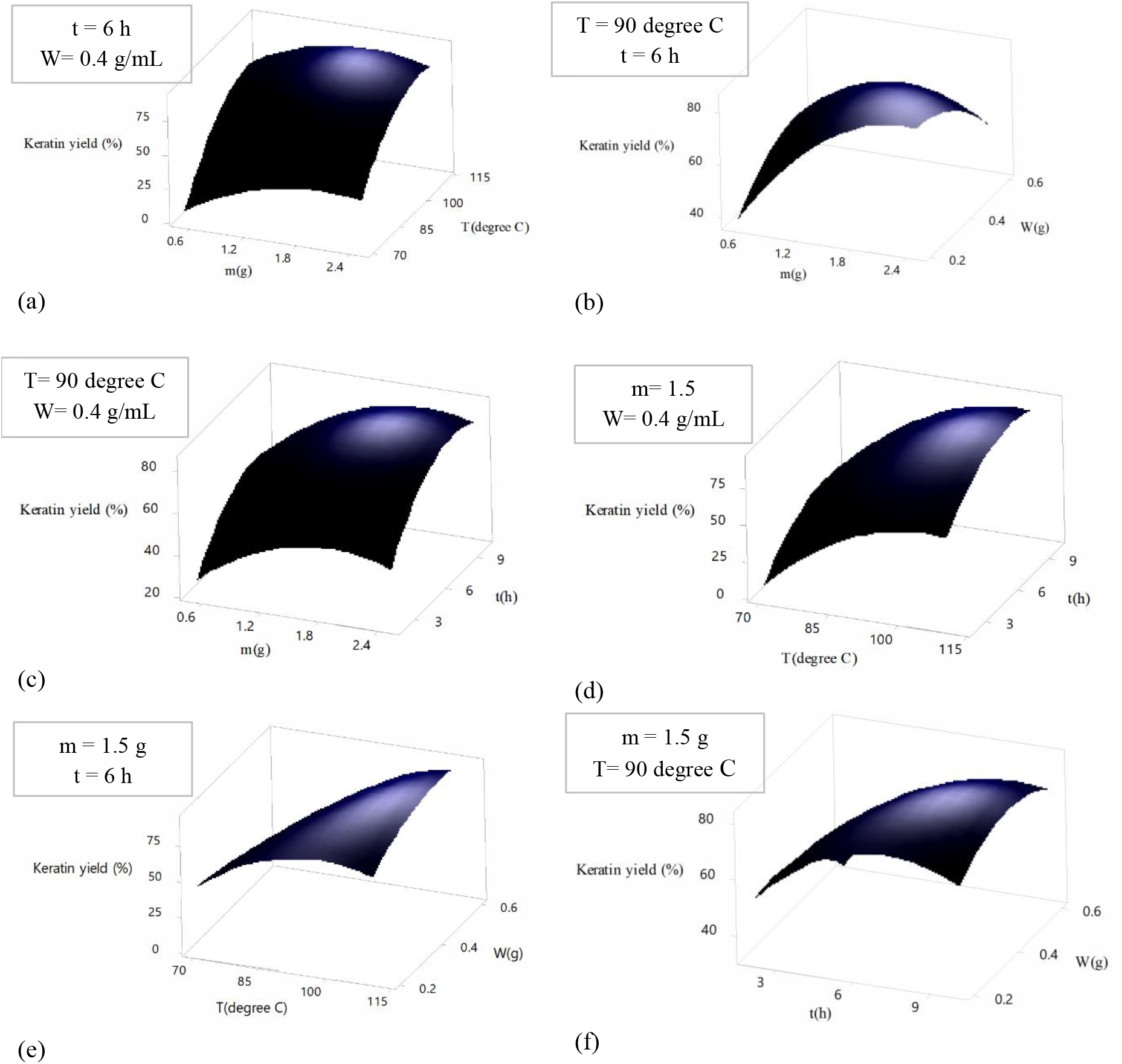
Surface plots showing the combined effects of processing variables on keratin yield, the legends highlight the values of the process variables held constant. Here T, t, W and m denote temperature (°C), time (h), wool load (g) and mass of L-cysteine in 20 mL of lactic acid (g), respectively. The legend shows the conditions at which the process variables were maintained constant.

### 3.3 Combined effects of variation in the process variables

Figure 2(a) shows the combined effect of changes in the mass of L-cysteine and changes in temperature on the keratin yield, at a dissolution time of 6 h and a coarse wool load of 0.4 g/mL. It can be observed that the combined increase in the mass of L-cysteine (from 0.5 g to 2.5g) and temperature (from 70 °C to 110 °C) leads to an overall increase in keratin yield (from 5.95 wt.% to ~75 wt.%). This is due to the combined favorable effects of the temperature on dissolution kinetics and L-cysteine on disulphide bonds cleavage. Crucially, although the combined effects of increasing temperature and mass of L-cysteine results in an overall increase in keratin yield, Figure 2(a) shows that at the highest temperature of 110 °C and highest mass of L-cysteine of 2.5 g, a marginal reduction in keratin yield is observed. This may be because of the associated thermal instability of the L-cysteine derived thiols at high temperatures.

Figure 2(b) shows the combined effect of changes in the mass of L-cysteine (from 0.5 g to ~2.5 g) and coarse wool load (0.2 g/mL to 0.6 g/mL) on the keratin yield at a dissolution time of 6 h and temperature of 90 °C. It is observed that the combined increase in the L-cysteine (from 0.5 g to 2.5g) and coarse wool load (from 0.2 g/mL to 0.6 g/mL) will lead to a net increase in the keratin yield (from 5.95 wt.% to ~75 wt.%). This observation highlights the dominance of the positive effect of higher masses of L-cysteine (section 3.2.1) compared to the negative effects of higher coarse wool load (section 3.2.4) and is consistent with discussions presented earlier above (section 3.1.2).

Figure 2(c) shows the combined effect of changes in the mass of L-cysteine and changes in time on the keratin yield, at a temperature of 90 °C and a coarse wool load of 0.4 g/mL. It can be observed that the combined increase in the mass of L-cysteine (from 0.5 g to 2.5 g) and time (from 2 h to 10 h) leads to an overall increase in keratin yield (from 5.95 wt. % to ~75 wt.%). This is due to the combined favorable effects of increasing time on dissolution kinetics and increasing mass of L-cysteine on disulfide cleavage stated earlier.

Figure 2(d) shows the combined effect of changes in temperature and changes in time on keratin yield at a L-cysteine mass of 1.5 g and a coarse wool load of 0.4 g/mL. As expected, it is observed that the combined effect of an increase in temperature (from 70 °C to 110 °C) and time (from 2 h to 10 h) leads to an increase in keratin yield (from 5.95 wt. % to ~75wt.%). This observation is a reflection of the favorable effects of increasing temperature and increasing time on wool dissolution kinetics, as discussed in section 3.2 above.

Figure 2 (e) shows the combined effects of changes in temperature and changes in coarse wool load on keratin yield when a L-cysteine mass and time is maintained at 1.5 g and 6 h, respectively. Increase in temperature (from 70 °C to 110 °C) and wool load (from 0.2 g/mL to 0.6 g/mL) resulted in an overall increase in keratin yield (from ~50 wt.% to ~75 wt.%). This observation indicates that the positive effect of increasing L-cysteine mass outweighs the negative impacts of increasing wool load (section 3.2). This observation is expected when the significance of the process variables discussed above (section 3.2.1) is acknowledged. Furthermore, Figure 2 (e) shows that if the wool load is maintained at 0.2 g/mL, higher temperatures will initially lead to higher keratin yields, with temperatures beyond an optimum value leading to a reduction in keratin yields. The decrease in keratin yield may be due to the degradation of keratin at higher temperatures [11].

Figure 2 (f) shows the combined effect of changes in time and changes in coarse wool load on keratin yield at a L-cysteine mass of 1.5 g and temperature of 90 °C. It is observed that the increase in time (from 2 h to 10 h) and the increase in wool load (from 0.2 g/mL to 0.6 g/mL) leads to an overall increase in keratin yield (from ~50 wt.% to 75 wt.%). This observation also indicates that the positive effect of an increase in time outweighs the negative effect of increasing wool load (section 3.2).

Figure 2 (f) also shows that if the wool load is maintained at 0.2 g/mL, increasing the time will initially favor higher keratin yields, then lead to poorer keratin yields at longer times. The decrease in keratin yield may be due to the degradation of keratin due to sustained heating of the keratin (at 90 °C) as the time increases significantly.

### 3.4 Determination of optimum conditions for keratin extraction

Employing the optimization module of the Minitab software, the empirical relation presented in Equation 4 was used to determine the conditions that maximize keratin yield. A keratin yield of 95 wt.% was predicted at 105 °C, 8 h, with 1.6 g of L-cysteine using 0.5 g of wool. To evaluate keratin yield at the suggested conditions, a set of experiments at the conditions were undertaken and the results presented in Table 4.

**Table 4.** Predicted and experimentally determined optimum keratin yield.

| $K_y$, wt.%<br>(Predicted) | $K_y$, wt.%<br>(Experiment) | Relative absolute deviation |
| --- | --- | --- |
| 95 | 93.77± 1.09 | 0.01 |

<sup>a</sup>Denotes relative absolute deviation; $K_y$ denotes keratin yield

Table 4 shows that the experimentally determined keratin yield at the optimum condition is 93.77 wt.%. This value is comparable to the predicted keratin yield of 95 wt.% with a relative absolute deviation (RAD) of 0.01 which is less than the minimum acceptable RAD of 5 %[37]. Indeed the experimental result confirms the validity of the experiment design method utilized and the sufficiency of the empirical model developed. Given that temperature was identified as the most important process variable in the keratin extraction from the waste coarse wool, we studied the potential effects of temperature on the structure of the resulting keratin. The physiochemical properties of the keratin were assessed at 70 °C, 90 °C and 110 °C using methods described in section 2.4 and the findings are presented in section 3.5.

### 3.5 Characterization of the obtained keratins

Here the effect of temperature on the keratin structure was assessed for extraction at 70 °C, 90 °C and 110 °C indicated as the low, middle, and high temperature values investigated in the present study. The keratin products generated at the low, middle, and high temperature conditions were designated as Keratin L, Keratin M and Keratin H, respectively. In all cases the dissolution time was maintained at 6 h. pictorial illustrations of Keratin L, Keratin M, and Keratin H are shown in Figures 3(a)-(c).

**Figure 3.**
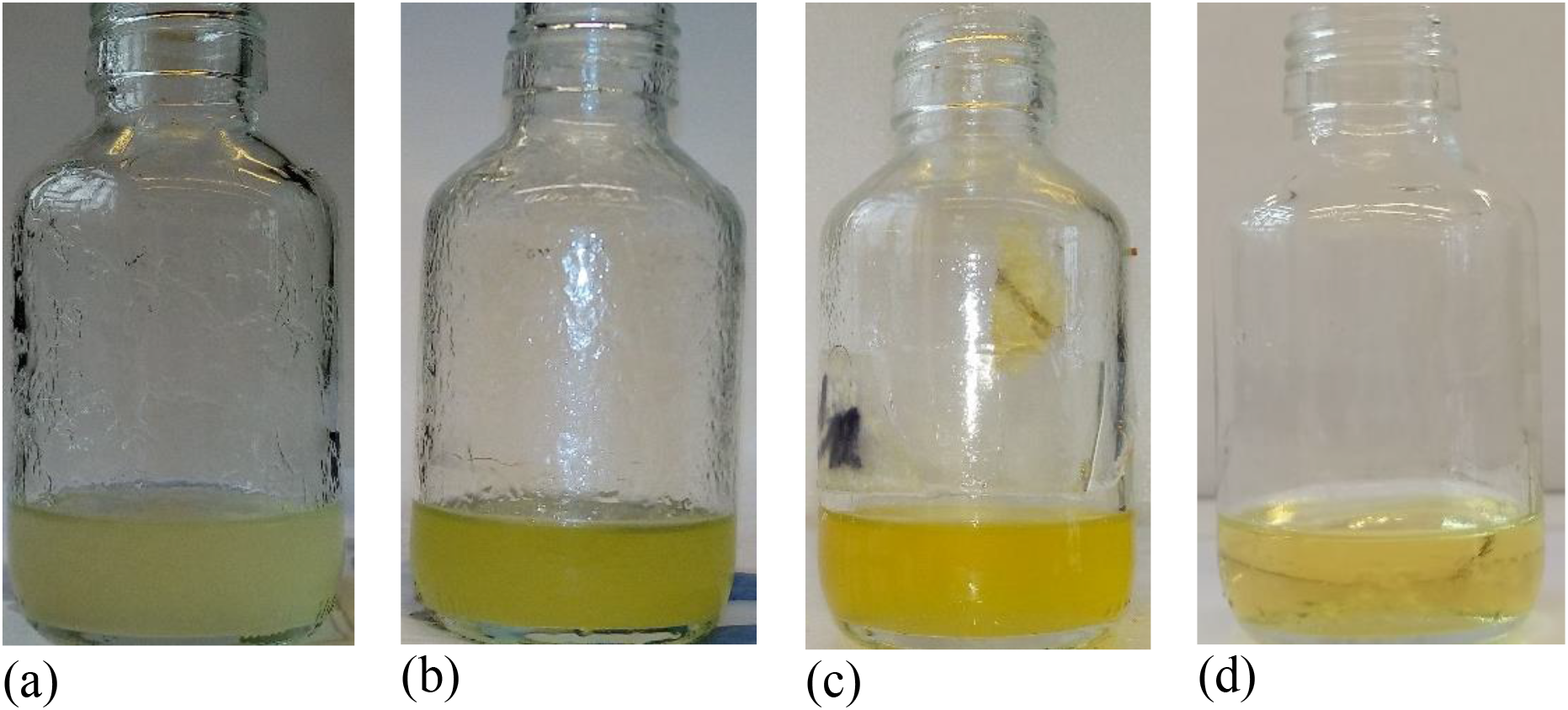
Keratin samples after 6 h processing at (a) 70 °C, (b) 90 °C and 110 °C (c) and the wool-free L-cysteine+lactic acid (DES) solvent at the upper-temperature limit of 110 °C, indicating that the yellowish color of the solvent is not related to the presence of the wool.

Figure 3 shows that the DES (Figure 3 d) has a yellow coloration, implying that the coloration in Figures 3(a)-(c) is due to the solvent. Visual investigation however shows that that the dissolution of the wool is enhanced as the temperature increases from 70 °C to 110 °C as expected. Employing the methods discussed in section 2 above, the keratin products generated at the different temperature conditions were characterized with the results presented and compared to the results from the analysis of coarse wool in Figure 4. Figure 4 shows that the FTIR spectra of coarse wool and keratin samples demonstrated similar spectra for all the samples indicating that the protein functional groups of keratin were retained after dissolution (Figure 4a). No significant difference was observed between FTIR spectra of keratin L, keratin M and keratin H. Peptide bonds of keratin were observed for the three regenerated keratin samples. Amide I characteristic band was observed for three samples at around 1640 cm^-1^ attributed to the C=O. Additionally, the characteristic band of amide II appeared at around 1530 cm^-1^ for the three keratin samples, which is assigned to stretching vibrations of the C–O and C–N [21, 38, 39]. Moreover, the presence of a peak at around 1040 cm^-1^ in keratin samples is due to the presence of cysteine-S-sulfonated residues in the recovered keratin [22, 40]. Hence, the result showed that keratin regeneration from wool using L-cysteine and lactic acid is a safe method for preserving the protein structure of keratin; however, choosing a proper experimental parameter, particularly dissolution temperature may be considered in large-scale applications.

**Figure 4.**
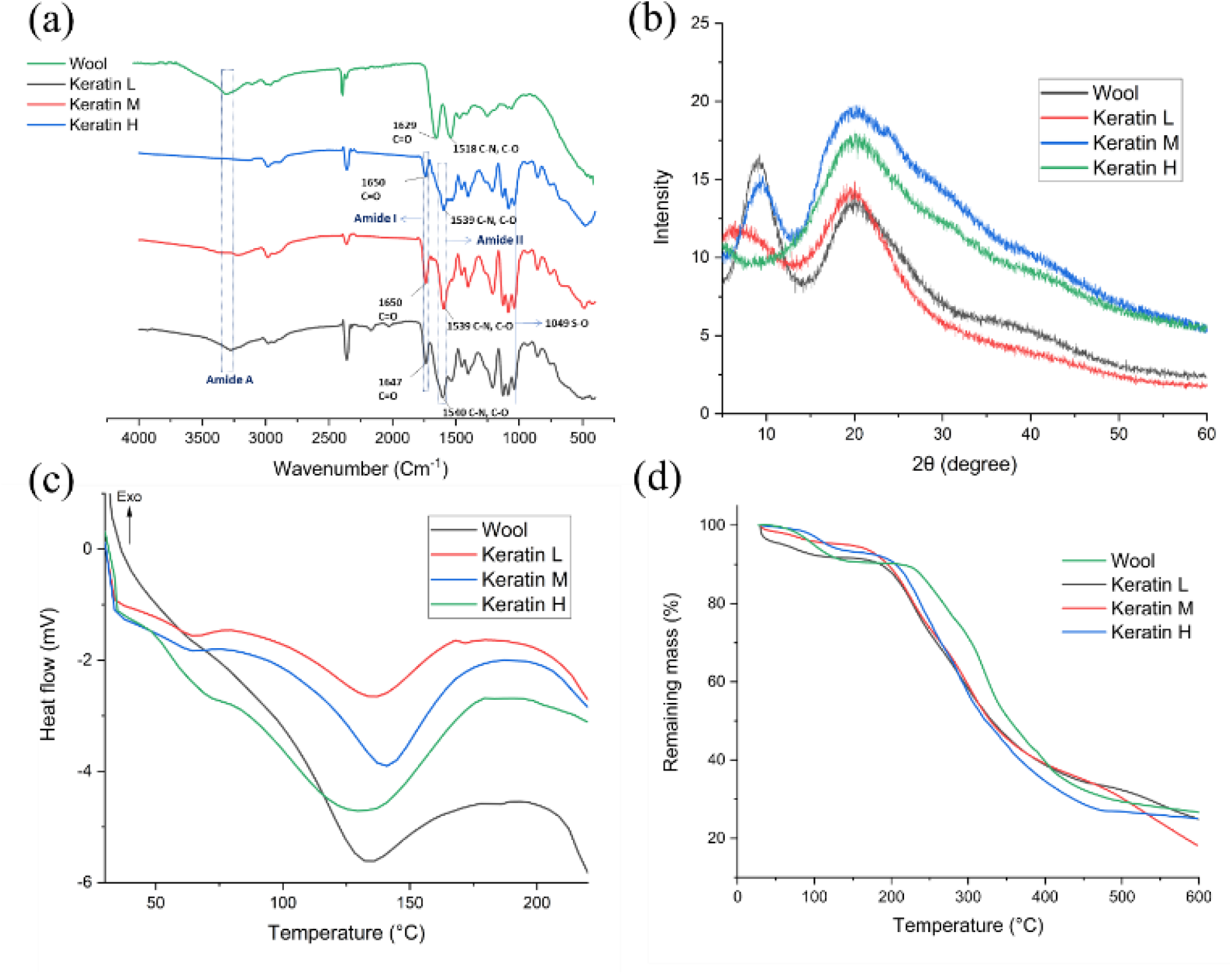
Physiochemical properties of regenerated keratin samples with various experimental parameters, (a) FTIR spectra, (b) XRD pattern, (c) DSC thermograms (30-240 °C, at heating rates: 10 °C min^-1^) and (d) TGA curves of keratin samples (30 to 600 °C at 10 °C min^-1^).

Moreover, the X-ray diffraction pattern of wool and three keratin samples were evaluated (Figure 4b). Characteristic bands around 10° and 20° were observed, aligning with the α-helix structure and β-sheet structure of keratin [22]. The results showed that the intensity of the β-sheet structure’s peak increased after the dissolution compared to the wool and further increased by increasing the dissolution temperature from 70 (keratin L) to 110 °C (Keratin H). Keratin M showed the highest peak intensity with a characteristic peak at 20°, indicating a higher β-sheet structure [22]. However, characteristic peaks at 10° completely disappeared in keratin H, indicating that the α-helix structure was destroyed during the dissolution process due to the high dissolution temperature (110 °C). Moreover, the crystallinity of the samples decreased as the dissolution temperature increased from (70-110 °C) since the crystallinity was shown to be 56.8, 28, 18.8 % for Keratin L, M, and H, respectively. The result showed that increasing the dissolution temperature, leads to the degradation of the α-helix structure and the loss of keratin crystallinity. Thermal properties of wool and dried keratin samples were evaluated by differential scanning calorimetry (DSC) and thermogravimetric analysis (TGA) (Figures 4 c, d). All samples showed an endothermic peak around 80-150 °C attributed to the water loss and an exothermic peak of 180-220 °C, which could be due to thermal denaturation of the α-helical crystallites [41, 42] and peptide bonds degradation [43] according to the DSC results (Figure 4 c). Moreover, the results demonstrated that the integration of the exothermic peak decreased after the dissolution from keratin L to Keratin H, and a broad peak observed for the keratin samples around 200 °C could be due to reducing the molecular weight of the keratin samples and the amount of α-helix structure. Similarly, the TGA result showed two-stage of degradation for the wool and keratin samples. Samples showed the first degradation stage at around 80-150 °C with a mass loss of 9.59 % for the wool sample and about 4 % for the keratin samples attributed to the samples’ water loss. Furthermore, the second degradation stage occurred between 200 to 500 °C with mass loss of around may be due to the degradation of keratin polypeptides. The maximum degradation temperatures (DTGmax) of the wool sample was 323 °C, and for keratin, samples were 304, 300, and 289 °C for keratin L, keratin M, keratin H, respectively. The results showed that the thermal stability of keratin decreased in comparison to the wool samples.

### 3.6 Notable considerations

The present study emphasizes that apart from the high yields of keratin generated from wool when the novel DES is employed, several issues that have so far limited the environmental sustainability of wool dissolution for keratin recovery are also circumvented. For instance, the use of peroxides, sulfites, dithiothreitol and thiols, recognized as toxic and unsustainable as well as the use of high temperature and pressure approaches characterized by high costs, are avoided[44–47]. Furthermore, although the use of enzymes for keratin extraction is recognized as environmentally benign, the long durations (months) required to achieve wool dissolution and the long enzyme production cycles, make the use of enzymes [48], impractical. The novel DES in the present study solves these issues mentioned above, since the wool dissolution process is fast (optimal yield of > 90% at 8 h) and the DES is environmentally benign. However, the present study acknowledges that important cost concerns that may limit the applicability of the use of the novel DES in wool dissolution in large-scale systems. This is because the l-cysteine (food grade US$ 20-US$ 35 per kg [49]) component of the DES has been reported to have a higher market price compared to the market prices costs of the more toxic alternatives such as 2-mercaptoethanol (US$ 2.4-US$ 2.6 per kg [50]). However, given that the l-cysteine serves as a hydrogen donor (i.e. l-cysteine is oxidized) to facilitate the cleavage of the disulfide bond, it suggests that the l-cysteine may be recovered via its oxidation. Indeed, the l-cysteine redox reaction reversibility was highlighted in the literature [51].

According to Figure 5, the oxidized l-cysteine can be ‘regenerated’ by acidification (i.e using a Brønsted-Lowry acid). Therefore, we hypothesize that the regeneration of l-cysteine can be achieved by introducing stoichiometrically determined amounts of lactic acid, thus retaining favorable environmental outcomes. We anticipate that such regeneration of l-cysteine will circumvent the economic concerns since the opportunity of l-cysteine recycle will exist. Investigations into the practicability of such l-cysteine recovery in continuous systems will therefore constitute the basis of future studies within the research group.

**Figure 5.**
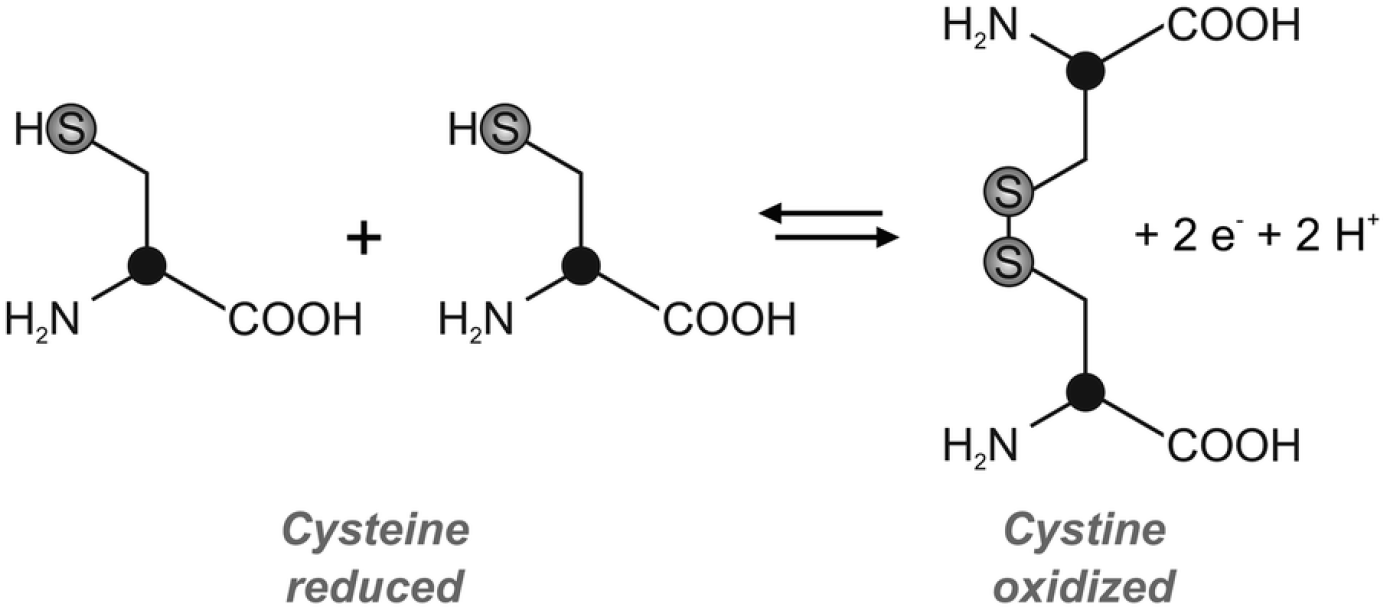
Redox reaction of l-cysteine [51].

## 4 Conclusion

In this work, the dissolution of waste coarse wool using a deep eutectic solvent (DES) mixture of L-cysteine and lactic acid has been investigated. It was demonstrated that the DES can efficiently facilitate the dissolution of waste coarse wool via the cleaving of disulphide bonds in wool. This work also showed that the DES facilitated an optimal keratin yield of 93.77 wt. % at the temperature, dissolution time, a mass of L-cysteine in 20 mL of lactic acid and coarse wool load in the solvent of 102 °C, 3 h, 2.5 g and 0.58 g, respectively. Notably, a further consideration of the process parameters showed that the temperature constituted the most impactful process parameter influencing keratin yeild. The impact of temperature (70 °C, 90 °C and 110 °C) on the keratin structure was therefore subsequently investigated. The protein structure of keratin and thermal stability were preserved in all regenerated keratin samples. However, the samples’ crystallinity decreased as the temperature increased from 70 °C (keratin L) to 110 °C (keratin H). The α-helix structure of the regenerated keratin samples’ were also observed to decrease as the temperature increased. Our results showed that the DES mixture of L-cysteine and lactic acid constitutes a green candidate for keratin recovery, since the recovery was achievable without sacrificing the keratin polypeptide’s structure.

## Author Contributions

Conceptualization, O.V.O. L.N. and A.S. methodology, O.V.O., H.J and P.H. software, O.V.O..; validation, M.H, A.S. writing—original draft preparation, O.V.O. H.J., P.H., L.N., H.A. and A.S. writing—review and editing O.V.O. H.J., P.H., L.N., H.A. and A.S

## Acknowledgements

The first author (O.V.O) gratefully acknowledges the financial support of Wallonia-Brussels International via the Wallonie-Bruxelles International (WBI) excellence Postdoctoral fellowship. H.J acknowledge Innoviris Brussels, Belgium (https://innoviris.brussels) under the project 2019 – BRIDGE – 4: RE4BRU for his PhD fellowship. The content is solely the responsibility of the authors and does not represent the official views of the above-mentioned fellowship agencies.

